# The *Campylobacter concisus* BisA protein plays a dual role: oxide-dependent anaerobic respiration and periplasmic methionine sulfoxide repair

**DOI:** 10.1101/2023.06.12.544688

**Authors:** Stéphane L. Benoit, Robert J. Maier

**Affiliations:** Department of Microbiology, Center for Metalloenzyme Studies, Georgia, 30602; Department of Microbiology, University of Georgia, Athens, Georgia, 30602

## Abstract

*Campylobacter concisus*, an emerging pathogen found throughout the human oral-gastrointestinal tract, is able to grow under microaerobic or anaerobic conditions; in the latter case, N-or S-oxides could be used as terminal electron acceptors (TEAs). Analysis of 23 genome sequences revealed the presence of multiple (at least two, and up to five) genes encoding for putative periplasmic N- or S-oxide reductases (N/SORs), all of which are predicted to harbor a molybdopterin (or tungstopterin)-*bis* guanine dinucleotide (Mo/W-*bis*PGD) cofactor. Various N- or S-oxides, including nicotinamide N-oxide (NANO), trimethylamine N-oxide (TMAO), biotin sulfoxide (BSO), dimethyl sulfoxide (DMSO) and methionine sulfoxide (MetO), significantly increased anaerobic growth in two *C. concisus* intestinal strains (13826 and 51562) but not in the *C. concisus* oral (type) strain 33237. A collection of mutants was generated to determine each N/SOR substrate specificity. Surprisingly, we found that disruption of a single gene, annotated as “*bisA*” (present in strains *Cc*13826 and *Cc*51562, but not in *Cc*33237) abolished all N/S-oxide-supported respiration. Furthermore, Δ*bisA* mutants showed increased sensitivity to oxidative stress and displayed cell envelope abnormalities, suggesting BisA plays a role in protein MetO repair. Indeed, purified recombinant *Cc*BisA was able to successfully repair MetO residues on a commercial protein (β−casein), as shown by mass spectrometry. Our results suggest that BisA plays a dual role in *C. concisus*, by allowing the pathogen to use N/S-oxides as TEAs, and by repairing periplasmic protein-bound MetO residues, therefore essentially being a periplasmic methionine sulfoxide reductase (Msr). This is the first report of a Mo/W-*bis*PGD-containing Msr enzyme in a pathogen.

**IMPORTANCE:** *C. concisus* is an excellent model organism to study respiration diversity, including anaerobic respiration of physiologically relevant N/S-oxides compounds, such as BSO, DMSO, MetO, NANO, and TMAO. All *C. concisus* strains harbor at least two, often three, and up to five genes encoding for putative periplasmic Mo/W-*bis*PGD-containing N/S-oxide reductases. The respective role (substrate specificity) of each enzyme was studied using a mutagenesis approach. One of the N/SOR enzymes, annotated as “BisA”, was found to be essential for anaerobic respiration of both N- and S-oxides. Additional phenotypes associated with disruption of the *bisA* gene included increased sensitivity toward oxidative stress and elongated cell morphology. Furthermore, a biochemical approach confirmed that BisA can repair protein-bound MetO residues. Hence, we propose that BisA plays a role as a periplasmic methionine sulfoxide reductase. This is the first report of a Mo/W-*bis*PGD-enzyme supporting both N-or S-oxide respiration and protein-bound MetO repair in a pathogen.

## INTRODUCTION

The Gram-negative bacterium *Campylobacter concisus* (1) belongs to the order Campylobacterales, a group that includes well-studied human pathogens such as *Helicobacter pylori* and *Campylobacter jejuni*. Although *C. concisus* has been found in saliva of healthy people, nevertheless it is commonly viewed as an emerging pathogen (2). Indeed, a growing body of evidence correlates the presence of the bacterium with various pathologies and diseases of the entire human oral-gastro-intestinal tract (OGIT), including (i) periodontitis, gingivitis and other dental diseases (3); (ii) Barrett’s esophagus (4); (iii) gastroenteritis and diarrhea (5–9); (iv) inflammatory bowel disease in adults (10, 11); (v) Crohn’s disease in children (12); (vi) microscopic colitis (13, 14). The distribution of *C. concisus* throughout the entire OGIT, composed of diverse environments in term of redox conditions and available nutrients, implies the presence of versatile respiratory pathways (15, 16). *C. concisus* can grow under both anaerobic or microaerobic conditions, however hydrogen (H_2_) is required for optimal growth (2, 15, 17–19). In addition to H_2_, putative electron donors include succinate, formate, malate, and reduced flavodoxin (15, 16, 20). *C. concisus* is expected to use a wide range of electron acceptors, including oxygen, fumarate, nitrate, nitrite, nitric oxide, nitrous oxide, tetrathionate, thiosulfate, as well as various N/S-oxides, the topic of the present study (15, 16).

Analysis of all *C. concisus* genomes available to date highlights the presence of several (*i.e*., at least two, often three or four, and sometimes up to five) genes encoding for putative <u>N</u>- or <u>S</u>-<u>o</u>xide reductases (hereafter referred to as “N/SORs”); all of which are predicted to harbor a molybdopterin (or tungstopterin)-*bis* guanine dinucleotide (Mo/W-*bis*PGD) cofactor. In comparison, the reference strain *C. jejuni* NCTC 11168, previously shown to reduce both DMSO and TMAO, contains only one related N/SOR protein complex (21–23). Recently, Yeow *et al.* found that the anaerobic growth of three *C. concisus* strains was significantly better when DMSO was added on plates, suggesting that this particular S-oxide can be used as alternative electron acceptor by *C. concisus* (16). However, DMSO reductase activity was not assigned to any specific gene or enzyme in any of these three strains (16). In the present study, we selected three unrelated *C. concisus* sequenced strains, *i.e.,* 13826, 33237, and 51562, to study N/S oxide respiration for the following reasons: (i) they are representative of the two main *C. concisus* genomospecies (GS), as both *Cc*51562 and *Cc*33237 belongto GS1, while *Cc*13826 belongs to GS2 (24, 25); (ii) they come from distinct human geographic niches, as both *Cc*13826 and *Cc*51562 were isolated from feces (6), whereas *Cc*33237 was isolated from the oral cavity (25); (iii) finally, they harbor very distinct sets of N/SOR genes, representative of *C. concisus* as a species.

## RESULTS

### Abundance of periplasmic Mo/W-*bis*PGD N- or S-oxide reductases in *C. concisus*

Genome analysis of 23 available *C. concisus* genomes (24–26) revealed the presence of multiple genes encoding for Mo/W-*bis*PGD-containing N/SOR catalytic subunits (**Table S1**), including in the three *C. concisus* strains used in this study. Indeed, strains 13826, 33237, and 51562 contain five, three, and two putative N/SOR genes, respectively (**Fig.1**, **Table S2**). Likewise, strains P26UCO-S2 and P2CDO4, previously shown to display DMSO-enhanced anaerobic growth (16) contain three and four putative N/SOR genes, respectively. Although there is limited homology between them (**Table S3**), nonetheless all *C. concisus* N/SOR protein sequences share three characteristics: (i) the presence of a N-terminus twin arginine translocation (Tat)-dependent peptide, indicative of periplasmic location (27), as predicted by both PRED-TAT (28) and SIGNAL P 6.0 (29) (**Fig. S1**); (ii) conserved motifs predicted to coordinate a Mo/W-*bis*PGD cofactor; (iii) conserved motifs predicted to bind four iron-four sulfur ([4Fe-4S]) clusters.

**Fig. 1.**
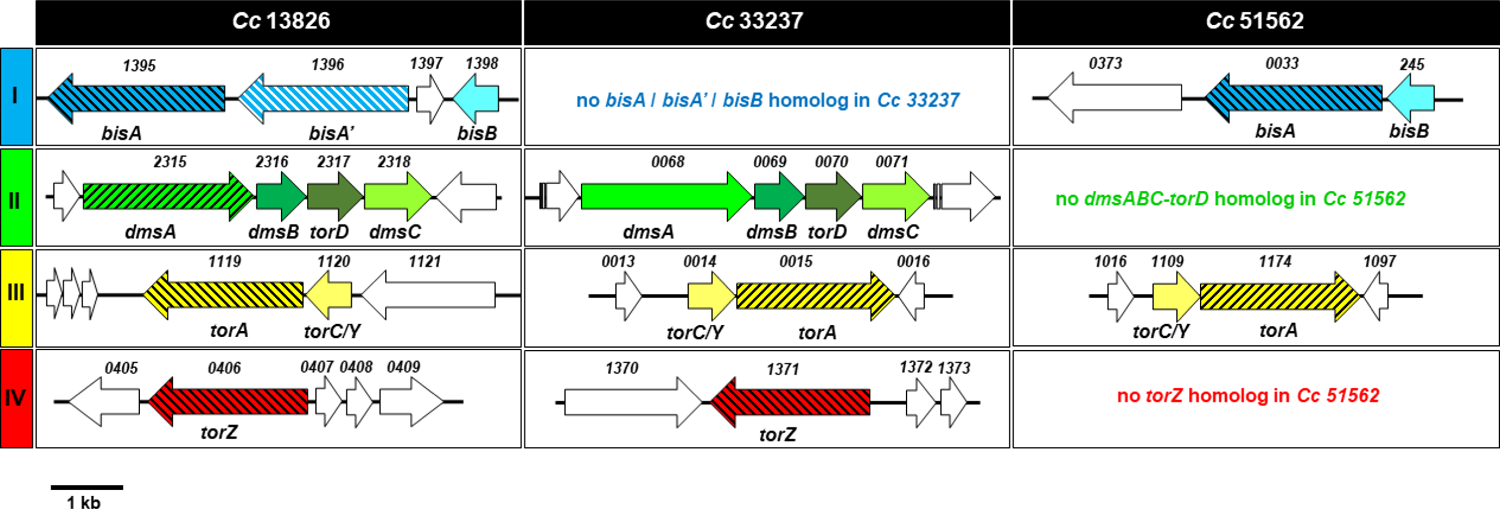
Location and organization of genes encoding for putative N- and S-oxide reductases in various *C. concisus* strains, and mutagenesis sites. The name of each *C. concisus* strain (13826, 33237 and 51562) is indicated above each column. Each gene is represented by an arrow. Colored arrows represent genes encoding for NSOR components and white arrows represent genes not relevant to the study. Putative gene names and locus tag numbers are indicated underneath and above each arrow, respectively. Gene annotations are according to JGI-IMG/M website (*<u>img.jgi.doe.gov</u>*). Genes targeted in this study (*bisA*, *bisA’*, *dmsA*, *torA*, *torZ*) are indicated by hatched arrows. An approximate scale is shown bottom left.

The three *C. concisus* strains studied herein (13826, 33237, and 51562) have only one N/SOR complex in common, the TorA-TorC/Y complex conserved in all *C. concisus* strains (**Table S1**). The *torA* gene encodes for a putative 813 amino-acid (aa)-long catalytic subunit, while the adjacent *torC/Y* gene encodes for a putative 189 aa-long, monoheme-containing cytochrome *c*, predicted to be periplasmic, based on the presence of a Sec-dependent signal peptide (28, 29). Another putative N/SOR gene, annotated as “*torZ*”, is usually found by itself (*i.e*., without any adjacent cytochrome-encoding gene). This gene, which encodes for a 817 aa-long periplasmic reductase, is present in almost all *C. concisus* strains (*e.g*., 21 out of 23 sequenced genomes, **Table S1**), including *Cc*13826 and *Cc*33237; however, *torZ* was not found in the genome of *Cc*51562.

Other putative N/SOR genes encountered in many (87%) *C. concisus* genomes include genes annotated as “*bisA*” and “*bisB*”; both are present in *Cc*13826 and *Cc*51562, however neither can be found in *Cc*33237 (**Table S1**). The *bisA* gene encodes for a putative 855 aa-long catalytic subunit, while the *bisB* gene encodes for a 189 aa-long putative monoheme type *c*-cytochrome. Both BisA and BisB are expected to be transported to the periplasm, *via* the Tat and Sec translocation system, respectively (28, 29) (**Fig. S3**). Among all putative N/SORs found in *C. concisus*, BisA is the catalytic subunit that shares the highest homology (65% identity, 78% similarity) with the *C. jejuni* Cj0264c protein found in strain NCTC 11168 (21) (**Table S2**).

In addition, one fourth of *C. concisus* genomes harbor the *dmsAB*-*torC*-*dmsC* operon, predicted to encode for a DmsABC-type heterotrimeric membrane-bound complex (**Fig. S3**) similar to that found in *E. coli* (30) and in select strains of *C. jejuni* (31, 32) (**Table S2**). This operon is present in strains *Cc*13826 and *Cc*33237, however it was not detected in the *Cc*51562 genome. Finally, *Cc*13826 contains yet another putative N/SOR gene: the 13826*_1396* gene, adjacent to *bisA* (*1395*) and *bisB* (*1398*) (**Fig. 1**), encodes for a 835 aa-long protein with high sequence homology to BisA (62% identity). The presence of this seemingly redundant gene, herein annotated as *bisA’*, is restricted to a few (∼22%) *C. concisus* strains (**Table S1**).

### Addition of various S- or N-oxides enhanced anaerobic respiration in the intestinal strains *Cc*13826 and *Cc*51562, but not in the oral strain *Cc*33237

*C. concisus* WT strains 13826, 33237, and 51562, were incubated in rich liquid broth supplemented with either BSO, DMSO, MetO, NANO, or TMAO (10 mM each), and their growth yield after 24 h was compared to that achieved in non-supplemented “plain” medium (**Fig. 2**). Supplementation of the medium with any of the five N/S oxides significantly increased the growth yield in both intestinal strains, *Cc*13826 and *Cc*51562, as determined by OD_600_ read (**Fig. 2A** and **Fig. 2C**) and CFU counts (**Fig. 3**). By contrast, no increase in growth was observed in presence of 10 mM methionine sulfone (MetO_2_), a compound structurally related to MetO but which cannot be reduced by methionine sulfoxide reductases (data not shown). Surprisingly, none of the five N- or S-oxides had any effect on the growth of the (oral) type strain *Cc*33237 (**Fig. 2B**), even when oxide substrates were added in excess (up to 50 mM, data not shown). These unexpected results suggest *Cc*33237 cannot use any of the N-or S-oxide compounds for anaerobic respiration, despite the presence of three putative N/SOR complexes (*i.e*., *dmsABC*, *torAC/Y* and *torZ*) in its genome. Finally, addition of nitrate and oxygen to the growth medium and the overhead space, respectively, resulted in significant growth increase (data not shown), suggesting that the failure to use N-or S-oxide compounds is not linked to a broader respiration problem in this particular strain.

**Fig. 2.**
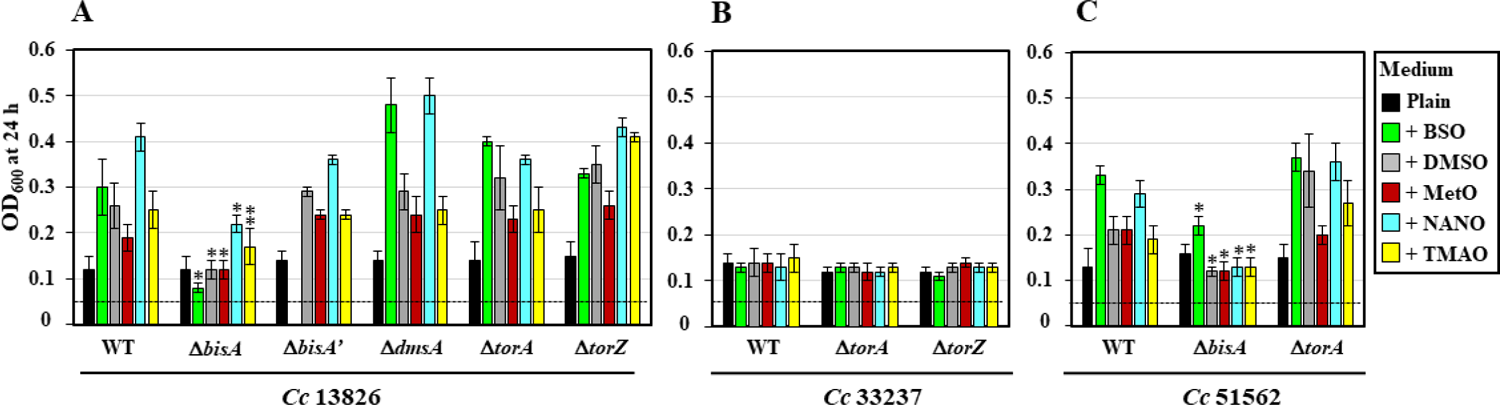
Growth yield of WT and mutant strains in presence of various N- or S-oxides. *C. concisus* WT strains 13826, 33237, 51562 and their isogenic N/SOR mutants were inoculated (OD_600_ of 0.05, dashed line) in plain growth medium, or in medium supplemented with 10 mM of either BSO, DMSO, MetO, NANO, or TMAO, as indicated by each colored square on the right. Cells were incubated under H_2_-enriched anaerobic atmosphere at 37°C, with permanent shaking (220 rpm), and OD_600_ was recorded after 24h. Results shown represent means and standard deviations from at least three biological replicates. A single asterisk above a bar indicates the bacterial growth yield (OD_600_) for the mutant is significantly lower compared to that of the WT strain, for the same (N/S oxide) condition (*P* < 0.01, Student’s *t*-test). Double asterisk, *P* < 0.02. The effect of BSO on the 13826 Δ*bisA’* mutant strain was not tested.

**Fig. 3.**
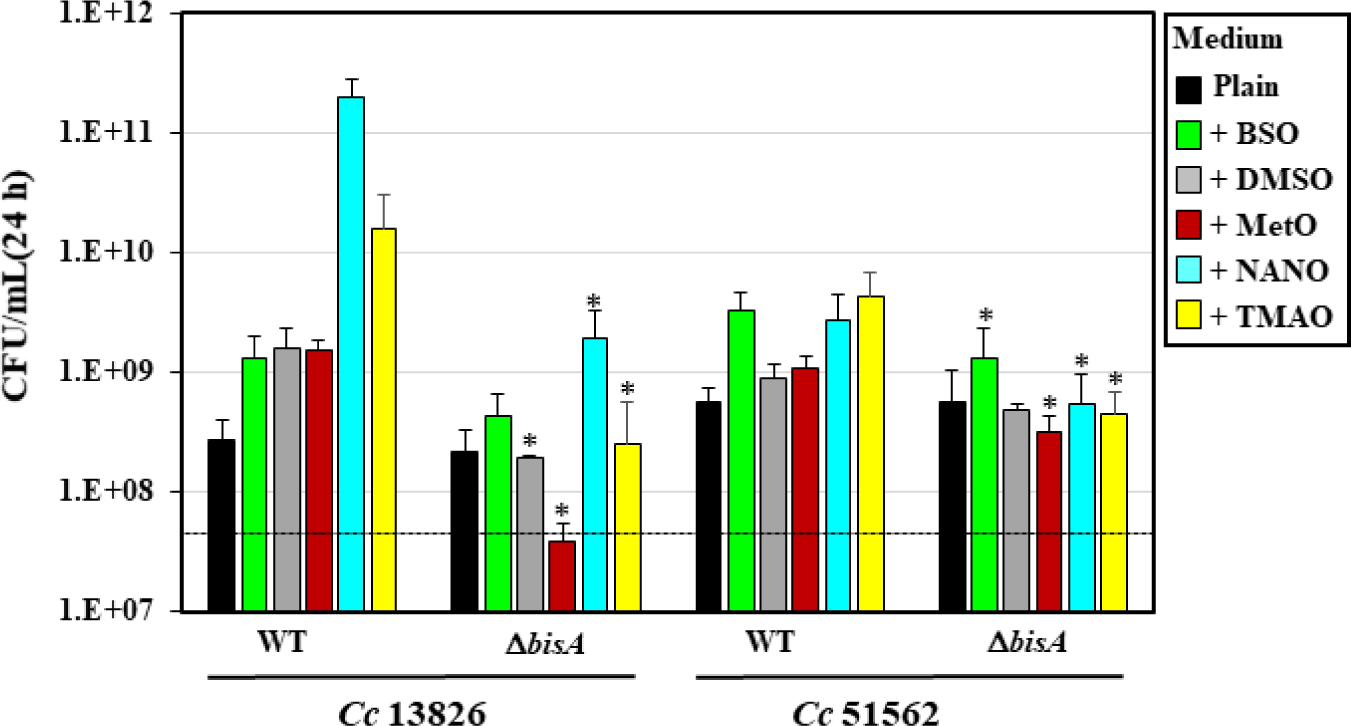
Quantitative growth yield of WT and Δ*bisA* mutant strains in presence of various N- or S-oxides. *C. concisus* WT 13826 and 51562, and Δ*bisA* mutant strains were inoculated (∼ 5 x 10^7^ CFUs/mL, dashed line) in plain growth medium, or in medium supplemented with 10 mM of either BSO, DMSO, MetO, NANO, or TMAO, as indicated by each colored square on the right. After 24 h, cells were serially diluted in PBS, spotted on BA plates in triplicate, and incubated under H_2_-enriched microaerobic conditions for two to three days, after which colony CFUs were counted. Results shown represent means and standard deviations of CFU/mL from three biological replicates, each with three technical replicates. Each color corresponds to a different growth medium (plain or 10 mM of each N/S oxide), as indicated on the right. The asterisk above each bar indicates the bacterial cell growth (CFU/mL) of the Δ*bisA* mutant under the indicated (N/S-oxide) condition is significantly lower compared to that of its isogenic WT strain grown under the same (N/S oxide) conditions (*P* < 0.01, Student’s *t*-test).

### *C. concisus* Δ*bisA* mutants are deficient in N/S-oxide-supported anaerobic growth

To determine which enzyme complex is responsible for N/S-oxide respiration in *C. concisus*, as well as to assign substrate specificity (*e.g*., N-versus S-oxide), nine different genes were inactivated among the three strains, as previously described (15) (**Fig. 1**). Each mutant was then compared to its respective parental strains for its ability to use any of the N/S oxide compounds as TEA during anaerobic growth (**Fig. 2**). In *Cc*13826, the *C. concisus* strain with the most diverse N/SOR gene content (*i.e*., five genes), mutagenesis of either *bisA’*, *dmsA*, *torA* or *torZ* did not lead to any growth deficiency. On the contrary, supplementation with either N/S-oxide resulted in significant anaerobic growth increase (compared to plain medium control) for either 13826 Δ*bisA’*, Δ*dmsA*, Δ*torA*, or Δ*torZ* mutant (**Fig. 2A**), suggesting that none of these four genes is involved in N/S-oxide reduction in *Cc*13826. Likewise, disruption of *torA* in strain *Cc*51562 did not lead to any deficiency in N-or S-oxide supported anaerobic growth; the 51562 Δ*torA* mutant grew as well as, and sometimes better than, the WT in presence of N/S-oxides (**Fig. 2C**).

By contrast, disruption of the *bisA* gene in strains *Cc*13826 (*i.e*., 13826_*1395*) and *Cc*51562 (*i.e*., 51562_*0033*) decreased or abolished N/S oxide-mediated growth stimuli, as shown by OD_600_ (**Fig. 2**) and live cells counts (**Fig. 3**). These results demonstrate that the BisA protein plays an essential role in N/S oxide respiration in both strains *Cc*13826 and *Cc*51562. Finally, despite the fact that no N/S oxide had any effect on the anaerobic growth of WT strain 33237 (see previous paragraph), two putative N/SOR genes (*i.e*., *torA* and *torZ*) were inactivated in this strain (**Fig. 1**), and their N-or S-oxide-dependent anaerobic growth was studied (**Fig. 2B**). As expected, none of the aforementioned compounds had any effect on the anaerobic growth of either mutant (**Fig. 2B**).

### *C. concisus* Δ*bisA* mutants have cell division defects

Upon routine microscopic observation of mutants, we noticed Δ*bisA* mutant cells had strikingly different morphology compared to their parental WT strain (**Fig. 4**). Indeed, bright field microscopy analysis revealed that both cells from strain 13826 Δ*bisA* (**Fig. 4B**) and 51562 Δ*bisA* (**Fig. 4D**) were significantly longer compared to their parental strains, WT strain 13826 (**Fig. 4A**) and 51562 (**Fig. 4C**), respectively. This unexpected phenotype is likely linked to the deficiency of the Msr activity of BisA (rather than its N/S-oxide respiratory role). We hypothesize that one or several periplasmic proteins involved in cell division in *C. concisus* are prone to methionine oxidation, and subsequent repair by BisA, as further discussed below.

**Fig. 4.**
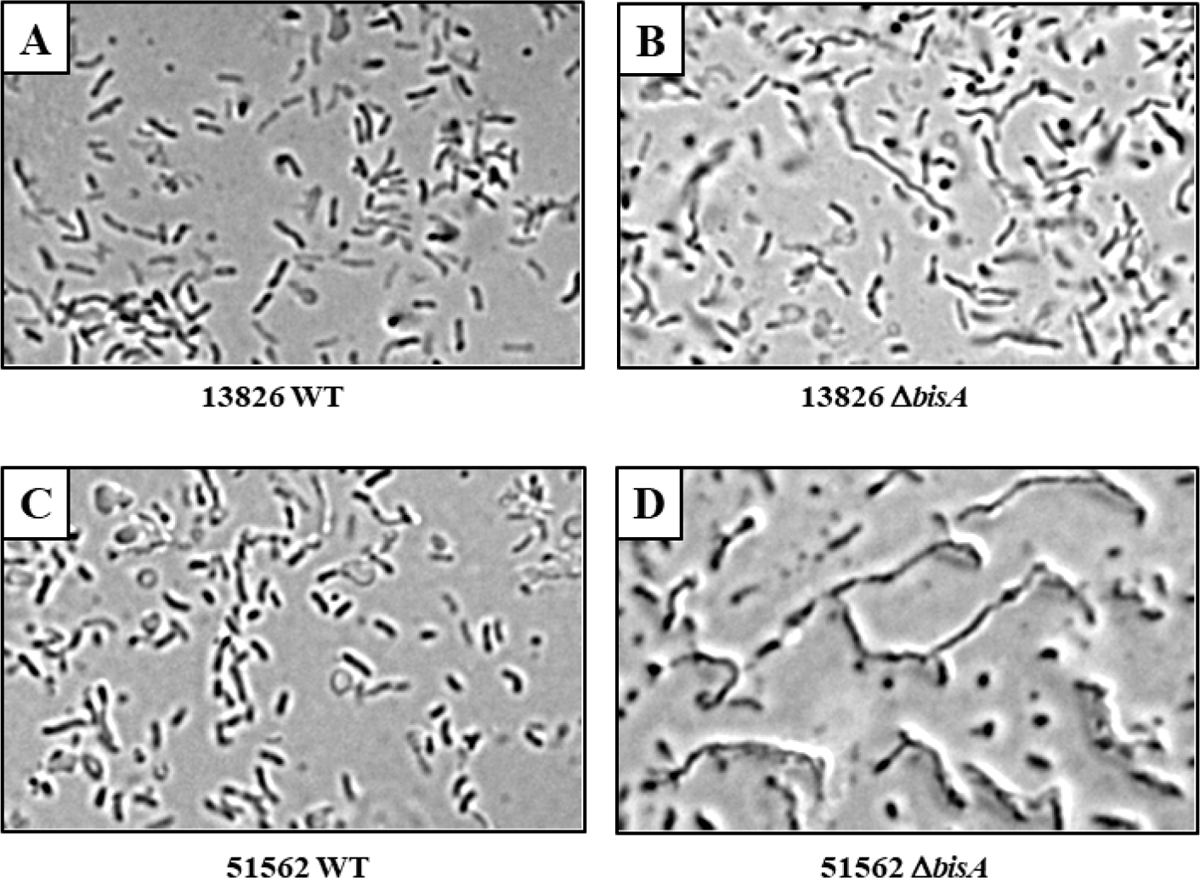
Bright field microscopy of *C. concisus* wild-type and Δ*bisA* mutant cells. *C. concisus* cells (13826 WT, 51562 WT and Δ*bisA* mutant strains) were grown for less than 24 h on BA plates under H_2_-enriched microaerobic conditions, resuspended in BHI broth, and examined with a bright field microscope (1000× magnification). Pictures have been digitally processed (brightness and contrast) for ease of interpretation.

### *C. concisus* Δ*bisA* mutants are more sensitive to oxidative stress

The inability to repair oxidized methionine residues usually correlates with increased sensitivity to reactive oxygen species (ROS), including superoxide anion (O ^-^.) and hypochlorous acid (HOCl) (33). Agar plate disk diffusion assays were used to compare the ability of *C. concisus* WT and Δ*bisA* mutant strains to combat methyl viologen (MV; a source of O_2_^-^.) or sodium hypochlorite (NaClO; a source of HOCl) (**Table 1**). Overall, WT strain *Cc*13826 appeared to be more sensitive (*i.e*., larger inhibition zones) than WT strain *Cc*51562 to both MV and NaClO. More importantly, both 13826 Δ*bisA* and 51562 Δ*bisA* single mutants were significantly more sensitive to both MV and HOCl compared to their respective parental strains (*P*< 0.001, Student’s *t*-test). The increased sensitivity of both Δ*bisA* mutants towards O_2_^-^. and HOCl, supports a role in MetO repair for the BisA periplasmic enzyme.

**Table 1.** Sensitivity of *C. concisus* WT and Δ*bisA* to methyl viologen and sodium hypochlorite.

| Strain | Diameter of inhibition zone (mm) |  |
| --- | --- | --- |
|  | Methyl viologen (100 mM) | Sodium hypochlorite (1.4 M) |
| 13826 (WT) | 23.5 ± 0.6 | 17.5 ± 0.6 |
| 13826 $\Delta bisA$ | 40.3 ± 0.5 | 23.8 ± 2.2 |
| 51562 (WT) | 17.0 ± 0.1 | 14.8 ± 1.1 |
| 51562 $\Delta bisA$ | 22.7 ± 0.6 | 18.0 ± 0.7 |

*C. concisus* strains 13826 and 51562 WT and isogenic $\Delta bisA$ mutant cells were grown on BA plates for 24 h before being resuspended in sterile PBS buffer to a final OD<sub>600</sub> of 2. Then 0.2 ml of cells were homogenously spread on top of 25-ml standardized BA plates. A sterile paper disk (7.5 mm diameter) was placed in the center of each plate, and 10 μL of 100 mM methyl viologen (MV) or 10 μL of 1.4 M sodium hypochlorite (NaClO) was added onto the disk. Cells were allowed to grow for 48 h, and the diameter (in mm) of the inhibition zone was measured. Results shown are means and standard deviations from three to five replicates. Inhibition zones (diameter) measured for each $\Delta bisA$ mutant were significantly larger than zones measured for each respective WT, for both MV and NaClO ( $P < 0.001$ , Student's *t*-test).

### In vitro repair of oxidized β-casein MetO residues by *Cc*BisA

Electron Spray Ionization-Mass Spectrometry (ESI-MS) was used to determine whether *Cc*BisA can repair oxidized β-casein *in vitro*. This approach has been successfully used to characterize several Msr enzymes, including *Rhodobacter sphaeroides* DorA and Msr, and *Saccharomyces cerevisiae* MsrB (34–36). Two variants of *C. concisus* BisA (native and hexahistidine-tagged) were expressed in *Escherichia coli* and purified to near homogeneity (**Fig. S4**). The N/S-oxide reductase activity of both purified proteins was spectrophotometrically assayed, using reduced benzyl viologen (BV) as electron donor, and either DMSO, MetO or TMAO as electron acceptor (data not shown).

Although both purified proteins showed measurable BV-dependent oxide reductase activity, including MetO reductase activity, the His-tagged version (*Cc*BisA-His_6_) was more active and therefore selected for the protein-bound MetO repair experiment. As mentioned above, the repair experiment involved the use of bovine β-casein, an intrinsically disordered protein that contains 6 Met residues readily oxidizable as either Met-R-O or Met-S-O epimer upon incubation with H_2_O_2_ (34). ESI-MS was used to measure the total mass of β-casein, before and after H_2_O_2_ treatment, and finally after incubation with *Cc*BisA-His_6_ (**Fig. 5**). Commercial β-casein consists of a mixture of genetic variants (34); in our case, up to eleven peaks were detected between 24,050 and 24,450 Da in the mixture (**Fig. 5A**). Prolonged treatment with H_2_O_2_ led to an overall mass increase of 96 Da for each variant (peak) corresponding to oxidation (*i.e*., + 16 Da) of each of the 6 Met residues, as previously reported (34) (**Fig. 5B**). Subsequent incubation of oxidized β-casein with *Cc*BisA-His_6_ in presence of dithionite-reduced BV (electron donor) resulted in a decrease in mass of approximately 16, 32, or 48 Da for most of the variants, corresponding to one, two, or even three MetO residues being repaired (*i.e*., recycled into Met), respectively (**Fig. 5C**). Similar changes (mass decrease) were not observed in the no-BisA control (oxidized β-casein with dithionite-reduced BV, data not shown), indicating BisA is responsible for the change in mass. Most (but not all) β-casein variants were repaired, however the fact that only one, two, or sometimes three MetO residues were repaired indicate that *Cc*BisA is stereospecific, *i.e*., it can only repair Met-R-O or Met-S-O epimers, but not both. Nevertheless, our results are in good agreement with previous reports. For instance, only one or two β-casein MetO residues were found to be repaired by *R. sphaeroides* DorA (34), and an average of 2.5 MetO per oxidized β-casein were reduced by *S. cerevisiae* MsrA (36). In conclusion, our proteomic results indicates *C. concisus* BisA is able to repair protein-bound MetO residues *in vitro*.

**Fig. 5.**
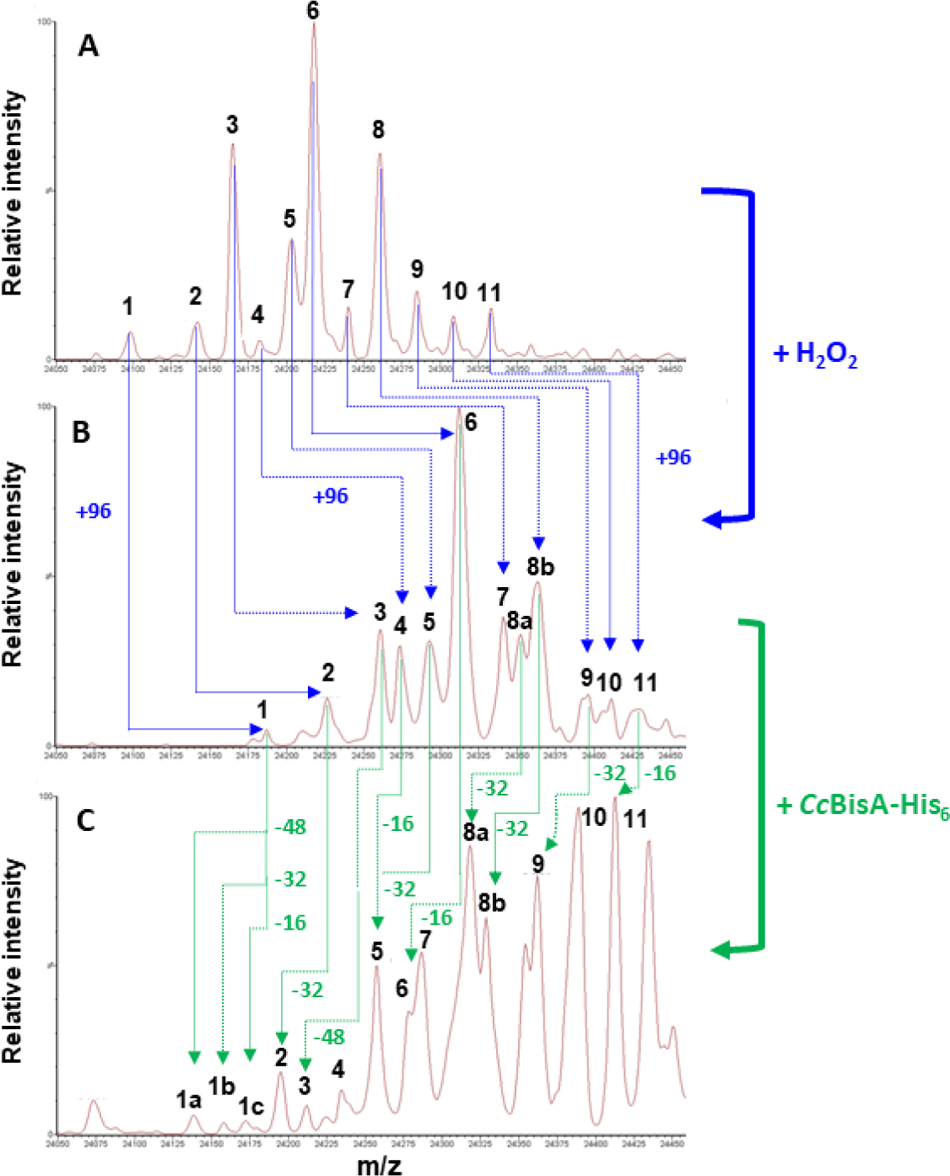
Oxidation of β-casein Met residues by H_2_O_2_ and reduction of MetO by *Cc*BisA(His)_6_, as analyzed by ESI-MS. (A) **Commercial bovine β-casein**. β-casein exists as a mixture of genetic variants; in this batch, eleven peaks are present between 24,050 and 24,450 Da. Peak mass (in Da): **1**. 24,098; **2**. 24,142; 3. 24165; **4**. 24182; **5**. 24023; **6.** 24,218; **7**. 24,240; **8**. 24,261; **9**. 24,285; **10**. 24,309; **11**. 24, 333. (B) **β-casein, following oxidation with 200 mM H_2_O_2_.** An increase of 96 (±4) Da (corresponding to six Met residues being oxidized) was observed for all peaks, as indicated with dotted blue arrows. Peak mass (in Da): **1**. 24,187; **2**. 24,226; **3**. 24,261; **4**. 24,274; **5**. 24,293; **6**. 24,312; **7**. 24,314; **8a**. 24,352; **8b**. 24,363; **9**. 24,396; **10**. 24,411; **11**. 24,429. (C) **Oxidized β-casein, after incubation with *Cc*BisA(His)_6_.** A decrease of either 16, or 32, or 48 (±2) Da (corresponding to one, two, or three MetO residues being repaired, respectively) was observed for most peaks, as indicated with dotted green arrows. Peak mass (in Da): **1a**. 24,138; **1b**. 24,156; **1c**. 24,172; **2**. 24,195; **3**. 24,212; **4**. 24, 235; **5**. 24,258; **6**. 24,277; **7**. 24,287; **8a**. 24,318; **8b**. 24,329; **9**. 24,362; **10**. 24,389; 11. 24,413.

## DISCUSSION

### Role of BisA in N/S oxide-supported anaerobic respiration

The ability to use alternative electron acceptors for respiration confers an advantage to the organisms that are capable to do so. For instance, both DMSO and TMAO have been known for a long time to be excellent electron acceptors for a variety of bacteria, including purple photosynthetic bacteria such as *R*. *sphaeroides* and *R. capsulatus* (37), γ-proteobacteria such as *Escherichia coli* (38) and related ε-proteobacteria such as *C. jejuni* (22, 23) and *Wolinella succinogenes* (39). Based on results presented herein and elsewhere (16), it is now clear that the list of DMSO-reducing bacteria should include *C. concisus*. We must however restrict our conclusion to the two intestinal strains used in our study (13826 and 51562) (24), since the oral strain *Cc*33237 (25) did not respond to the DMSO supplementation treatment. Besides DMSO, four other N/S oxides, BSO, MetO, NANO and TMAO, were shown to significantly bolster the growth of both *Cc*13826 and *Cc*51562, but not that of *Cc*33237. Although puzzling at first, the apparent inability of strain *Cc*33237 to respire any N/S-oxide compounds appears to correlate with the absence of a *bisA* homolog in the genome of this strain, hence reinforcing the role of BisA as the main N/SOR enzyme in *C. concisus*. To confirm this hypothesis, it would be of interest to determine whether other *bisA*-lacking *C. concisus* strains, including the previously studied P10CDO-S2 or H1O1(40), fail to respond to the N/S-oxide supplementation treatment. Alternatively, introducing and expressing *bisA* in strain *Cc*33237 would provide us with useful information. However, such heterologous expression has never been done in *C. concisus*, and it is anticipated to be challenging due to the paucity of genetic tools.

Unexpectedly, disruption of each of the four other putative N/SOR genes encountered in *C. concisus* (*i.e*., *bisA’*, *dmsA*, *torA*, *torZ*) did not lead to any detectable phenotype, leaving the respective role of each of these enzymes unanswered for now. In this context, previous studies conducted in the related ε-proteobacterium *C. jejuni* can be informative, although there are significant differences between N/SOR homologs found in the two *Campylobacter* species. Indeed, most *C. jejuni* strains, (*e.g*., NCTC 11168) contain only one N/SOR protein complex (Cj0264/0265 in strain NCTC11168) (21–23). In addition, a few *C. jejuni* strains (*e.g*., *Cj*81-176) possess a second N/SOR complex (31, 32) with significant homology to the DmsABC-type of DMSO reductase found in *E. coli* (30) and in some *C. concisus* strains (*e.g*., *Cc*13826 and *Cc*33237). Though *Cj* 81-176 Δ*dmsA* mutants have been found to display significant mouse colonization defects, suggesting *dmsA* plays a critical role *in vivo* (31), the DmsA enzyme has not been biochemically characterized and its preferred substrate is not known. Hence the physiological role of DmsA(BC) paralogs remains to be determined in *Campylobacter spp.* Regarding the Cj0264/0265 complex, the WT strain NCTC 11168 was shown to metabolize DMSO and TMAO (into DMS and TMA, respectively), a property lost in the *cj0264* mutant (23). Furthermore, both DMSO and TMAO were shown to serve as alternative electron acceptors by the WT, but not by the *cj0264* mutant (22, 23). Cj0264 is expected to get electrons from a monoheme *c*-type soluble cytochrome encoded by the adjacent *cj0265* gene, which is highly conserved in all *C. jejuni* strains (22, 41); Cj0265 itself is expected to receive electrons from the QcrABC membrane-bound menaquinol-cytochrome *c* reductase complex (42). In *C. concisus*, the concomitant presence of *qcrABC*, *bisA*, and *bisB* genes (the latter encoding for a monoheme *c*-type soluble cytochrome, highly similar to Cj0265) suggests a topological model and an electron transfer route similar to that described for *C. jejuni* Cj0264/264 (42) *i.e*., QcrC→BisB→BisA (**Fig. S3**).

### Role of BisA in periplasmic methionine sulfoxide repair

Based on various phenotypes associated with the disruption of *bisA* in both *Cc*13826 and *Cc*51562, we conclude that BisA plays a role as periplasmic Msr, able to repair both free MetO residues and protein-bound MetO. The first conclusion (repair of periplasmic free MetO) is based on the observation that free MetO can be used as respiratory substrate by BisA in the periplasm, *i.e*., MetO recycling into Met would be a direct consequence of the respiration process. Future studies using pure (BisA) enzyme will be needed to determine substrate stereospecificity (for instance, Met-*R*-O vs Met-*S*-O), as well as affinity constants and additional kinetic parameters. The second conclusion (repair of periplasmic protein-bound MetO) is based on the following results. Firstly, we found that the purified recombinant BisA-His_6_ enzyme was able to repair between one and three MetO residues out of six MetO residues found on oxidized β-casein, as shown by ESI-MS; these results are in agreement with that of previous studies using the same target protein (oxidized β-casein) and Msr-type enzymes such as *R. sphaeroides* DorA and *S. cerevisiae* MsrA (34, 36). Secondly, Δ*bisA* mutant cells failed to properly divide, suggesting BisA-mediated MetO repair is crucial for cell division in *C. concisus*; fully periplasmic, or membrane bound periplasmic-oriented Met-rich proteins that could account for such phenotype include two Met-rich proteins, annotated as “septum formation initiator” (6.3% Met content) and “cell division protein” (5.3% Met content); for reference, the average protein Met content in *C. concisus* is 2.4%. However, both the oxidation of these two periplasmic Met-rich proteins and their putative repair by BisA will need to be experimentally proven, using pure components and ESI-MS. Lastly, we observed an increased sensitivity of Δ*bisA* mutants toward oxidative stress, a hallmark of *msr* mutants in many eukaryotic and prokaryotic organisms (33), including *Saccharomyces cerevisiae* (43), *E. coli* (44), *H. pylori* (45) and *C. jejuni*, (46, 47).

*C. concisus* contains an additional periplasmic Msr enzyme, however it is not of the MsrPQ-type originally described in *E. coli* (48) and widely encountered in many *Campylobacter sp*., including in *C. jejuni* (42, 49). Instead, *C. concisus* contains a homolog of the multifunctional MsrAB-type fusion protein, similar to that found in *H. pylori* (45), but with one notable difference: *Hp*MsrAB is cytoplasmic, whereas *Cc*MsrAB is predicted to be periplasmic (29). Surprisingly, genome sequence analysis suggests there is no cytoplasmic Msr in *C. concisus*. This finding raises the question of the fate of cytoplasmic oxidized proteins, especially in the catalase-negative organism that is *C. concisus*. In the absence of a dedicated cytoplasmic MetO repair pathway, a non-specific quenching mechanism could be involved, similar to that previously uncovered in *H. pylori* (50) or in other organisms, including in eukaryotes (51, 52).

### Some BisA homologs play similar dual roles in other bacteria

Two periplasmic molybdoenzymes with significant homology to *Cc*BisA have been shown to play a similar dual role (N/S-oxide respiration and MetO repair) in two unrelated organisms. In *Haemophilus influenzae*, a protein annotated as “MtsZ” can use BSO and MetO, and repair free MetO (53). However, purified *Hi*MtsZ could not repair calmodulin-bound MetO, suggesting that the primary role of *Hi*MtsZ is respiratory, rather than MetO-protein repair (53). This is a key difference with *Cc*BisA, since a repair role of both free MetO and protein-bound MetO is proposed here. In the photosynthetic bacterium *R. sphaeroides*, a protein named “DorA” also plays a role in both MetO respiration and repair (34). Indeed, *Rs*DorA was shown to repair two physiologically relevant periplasmic oxidized Met-rich proteins, including the copper chaperone PCu_A_C (34). Interestingly, a periplasmic Met-rich (*i.e*., 6.4% Met) PCu_A_C homolog is present in *C. concisus*: it could be a preferred target for *Cc*BisA. There are differences though between *Cc*BisA and *Rs*DorA: *Cc*BisA is a soluble protein, predicted to be associated with only one protein, the soluble periplasmic cytochrome *c* annotated here as *Cc*BisB (**Fig. S3**), whereas *Rs*DorA is associated with two membrane subunits, DorB and DorC (34, 54), making the *R. sphaeroides* DorABC more similar to a DmsABC-type complex described above.

In summary, it appears that, among the many putative N/SORs found in *C. concisus*, only one, annotated as BisA, is actually involved in N/S-oxide-supported respiration. Additionally, *Cc*BisA is expected to repair protein-bound MetO in the periplasm. This dual role might be surprising, especially given the fact that *C. concisus* has another periplasmic MsrAB enzyme. But one has to remember that the average genome size of *Campylobacter sp.*, including *C. concisus* (*i.e*., 1.8-2.1 Mb), is significantly smaller compared to that of other groups of commensal or pathogenic bacteria living in similar environmental niches, including Staphylococci (∼3 Mb), Bacteroides (∼5-7 Mb), or Enterobacteriaceae such as *E. coli* and *S. enterica* (∼5 Mb). Thus, to adapt to and thrive in the extremely diverse environment that is the human OGIT, *C. concisus* has to rely both on versatile respiratory pathways and multifunctional enzymes, as exemplified herein by BisA, a bifunctional enzyme capable of both N/S-oxide respiration and protein-bound methionine sulfoxide repair.

## MATERIALS AND METHODS

### Bacterial strains and plasmids

*C. concisus* strains used in this study are listed in **Table S4**. The three *C. concisus* parental strains (13826, 33237, and 51562) were purchased from ATCC (Manassas, VA). Genomic DNA from each *C. concisus* parental strain was used as template for PCR to construct each strain-specific mutant. All PCR products were sequenced at ETON.

### Genome sequence analysis

Gene and protein sequences of *C. concisus* strains 13826 (also known as BAA-1457), 33237, and 51562 were obtained from the integrated microbial genomes (IMG) website of the Joint genome Institute (https://img.jgi.doe.gov), Biocyc (https://biocyc.org), and Genbank (www.ncbi.nlm.nih.gov), and Uniprot (www.uniprot.org).

Additional genome sequences were accessible via the FigShare link described in (26). Twin arginine translocation (TAT) sequences were predicted using PRED-TAT (28) and SIGNAL P 6.0 (29). Multiple protein sequence alignment and phylogenetic tree were made using Clustal Omega (https://www.ebi.ac.uk/Tools/msa/clustalo/).

### Chemicals

BSO (Biotin sulfoxide, #B389050) was from Toronto Research Chemicals, Toronto, ON, Canada. DMSO (Dimethyl Sulfoxide, #BP231) was from Fisher Scientific. MetO (L-methionine sulfoxide, #M1126), MetO_2_ (L-methionine sulfone, #M0876), NANO (Nicotinamide N-oxide, #M-3258), TMAO (Trimethylamine N-oxide, #317594), NaClO (Sodium hypochlorite #425044), and BV (Benzyl Viologen, #B8133) were all purchased from Sigma-Aldrich (Saint Louis, MO).

### Growth conditions

*Campylobacter concisus* was routinely grown on Brucella agar (Becton Dickinson, Sparks, MD) plates supplemented with 10% defibrinated sheep blood (Hemostat, Dixon, CA) (BA plates). Chloramphenicol (Cm, 8 to 25 μg/ml) was added as needed. BA plates were incubated at 37°C under H_2_-enriched microaerobic conditions, consisting of sealed pouches filled with anaerobic mix, a commercial gas mixture containing 10% H_2_, 5% CO_2_ and 85% N_2_ (Airgas, Athens, GA) and some residual air. For liquid cultures, 165-ml sealed bottles were filled with 2 to 10 mL of 1.25 X concentrated Brain-Heart Infusion (BHI, Becton Dickinson), autoclaved and then supplemented with 0.2 to 1 mL (10% final) of fetal calf serum (FCS, Gibco Thermo Fisher); and 0.2-1 mL (10% final) of either sterile deionized water (plain, control) or 100 mM BSO, DMSO, MetO, MetO_2_, NANO, or TMAO (each at a final concentration of 10 mM). Cells were grown under H_2_-enriched anaerobic conditions, as follows. First, bottles (overhead space) were sparged with N_2_ for 10 min, and then with anaerobic mix (see above) for 10 min. Additional H_2_ (10%) was added, to achieve 20% H_2_ partial pressure. The inoculum was prepared as follows: *C. concisus* cells grown on BA plates for less than 24 h were harvested, resuspended in BHI-FCS and standardized to the same OD_600_ before being inoculated. The starting OD_600_ was between 0.04 to 0.05, corresponding to approximately 5 x 10^7^ to 8 x 10^7^ cell forming units (CFU) per mL, respectively (15). Cells were grown (triplicate for each strain and condition) for 24 h at 37°C under vigorous shaking (200 rpm). Growth yield was estimated by both measuring OD_600_ and counting CFUs. For this purpose, samples from each bottle were serially diluted in PBS (up to 10^-8^), and 5 μL of each dilution was spotted in triplicate on BA plates. CFU were counted after 48 to 72 h of incubation under H_2_-enriched microaerobic conditions. Results shown are the mean and standard deviation (SD) of at least triplicate biological replicates, each with technical triplicate.

### Construction of *C. concisus* mutants

Each mutant was constructed by following a 3-step strategy, as previously described (15). Briefly, DNA sequences (400 bp- to 850 bp-long) flanking each target gene (*i.e.*, *bisA*, *bisA’*, *dmsA*, *torA*, *torZ*) were PCR-amplified, using genomic DNA from each *C. concisus* parental strain (*e.g*., 13826, 33237, 51562, see **Table S4**) and specific primers for each targeted gene (**Table S5**). Each set of two PCR products was combined with a 740 bp-long DNA sequence containing a *cat* (chloramphenicol resistance) cassette that has its own promoter {Wang, 1990 #116}, and the final PCR step yielded a product containing both flanking sequences with the *cat* cassette in between (more detailed information on the construction of each mutant is provided in Supplementary data). In the second step, each tripartite PCR product was purified and methylated, by incubating approximately 25 μg of DNA with 150-250 μg of (cell-free extract) total protein from *C. concisus* for 2 h at 37°C in presence of 0.4 mM S-Adenosylmethionine (SAM, New England Biolabs, Ipswich, MA). After methylation, each PCR product was purified again (Qiaquick purification kit, Qiagen, Valencia, CA), and the methylated DNA (1-5 μg) was introduced into *C. concisus*, by using natural transformation or electroporation (BTX Transporator Plus, 2,500 V/pulse). Transformed cells were first plated on BA plates and incubated (H_2_-enriched microaerobic conditions, see above) for 8-12 h before being transferred onto BA supplemented with 8-10 μg/mL Cm. Colonies appeared after 3 to 5 days. The concomitant deletion of the gene of interest and the insertion of *cat* was confirmed by PCR, using genomic DNA from mutants as template and appropriate primers.

### Microscopy analysis

*C. concisus* cells (13826 WT, 51562 WT and Δ*bisA* mutant strains) were grown for less than 24 h on BA plates under H_2_-enriched microaerobic conditions (see above), resuspended in BHI broth, and examined with a bright field microscope (Nikon Eclipse Ni). Digital images were obtained at 1000× magnification using a Nikon cool snap camera.

### N/S-oxide reductase enzyme assays

BisA reductase activity was measured as previously described (34, 35) with modification. Briefly, Britton–Robinson buffer (55), pH 6.0 was sparged for 20 min with N_2_ to create semi-anaerobic conditions. The N_2_-infused buffer was then mixed with 0.2 mM of Benzyl Viologen (BV, electron donor) and 5 to 10 mM of either DMSO, MetO or TMAO (N/S oxide electron acceptor) in a cuvette fitted with a silicone seal to prevent gas exchange (1 mL final volume). A few microliters of freshly prepared sodium dithionite (Sigma) solution were injected into the cuvette to reduce BV, until the absorbance at 600 nm was stable and between 1 and 1.5 units. The reaction was initiated by addition of 0.2 to 1 μM of purified *Cc*BisA or *Cc*BisA-His_6_. Oxidation of BV (*i.e*., A_600_ decrease) was followed for 1 min. Rates of N/S oxide reductase activity were calculated using a molar extinction coefficient (ε_600_) of 10,400 M^−1^·cm^−1^, with 2 moles of BV oxidized for 1 mole of reduced oxide substrate.

### ESI-MS analysis of β-casein oxidation and subsequent repair by *Cc*BisA

The protein-bound MetO repair ability of CcBisA was assessed using commercial bovine β-casein as target, and dithionite-reduced BV as electron donor, as previously described by Tarrago and collaborators, with modifications (34). Briefly, 100 μM of bovine β-casein (Sigma # C-6905) was oxidized overnight at room temperature in 10 mL PBS (pH 7.5) with 200 mM H_2_O_2_. The protein was concentrated and the oxidant was removed by using an Amicon concentrator with a MWCO of 10 kDa, followed by a PBS wash and a second concentration step. Finally, β-casein was resuspended in 1 mL of PBS, and the protein concentration was determined with the BCA protein assay kit. Some oxidized β-casein was kept aside for further analysis. For repair of oxidized β-casein by BisA, N_2_-sparged Britton-Robinson buffer, pH 6.0, was mixed with 100 μM β-casein, 1 μM of purified CcBisA-His)_6_ and 0.2 mM dithionite-reduced BV. Repair was conducted at 37°C for approximately 2 h, with 1.6 mM dithionite injected into the solution every 20 min, as previously described (34). The repair mix was frozen and stored at −80°C before being sent for analysis, along with appropriate controls (unoxidized β-casein; oxidized β-casein; and BisA-free, oxidized β-casein with BV and dithionite). The protein (β-casein) was purified by Liquid Chromatography (LC) and then analyzed by Electron Spray Ionization (53)-Mass Spectrometry (MS) on a Bruker Esquire 3000 plus ion trap mass spectrometer, at the Proteomics and Mass Spectrometry facility (PAMS), University of Georgia, Athens, GA.

## ACKNOWLEDGMENTS

This work was supported by the University of Georgia Foundation (RJM).

## Author contribution

SLB designed and conducted all experiments, and wrote the manuscript. RJM reviewed the manuscript. The authors wish to thank Lionel Tarrago (CNRS, Aix Marseille University, France) and Dennis Phillips (UGA PAMS) for help with this project.

